# Enhanced Brain Imaging Genetics in UK Biobank

**DOI:** 10.1101/2020.07.27.223545

**Authors:** Stephen M Smith, Gwenaëlle Douaud, Winfield Chen, Taylor Hanayik, Fidel Alfaro-Almagro, Kevin Sharp, Lloyd T Elliott

## Abstract

UK Biobank is a major prospective epidemiological study that is carrying out detailed multimodal brain imaging on 100,000 participants, and includes genetics and ongoing health outcomes. As a step forwards in understanding genetic influence on brain structure and function, in 2018 we published genome-wide associations of 3,144 brain imaging-derived phenotypes, with a discovery sample of 8,428 UKB subjects. Here we present a new open resource of GWAS summary statistics, resulting from a greatly expanded set of genetic associations with brain phenotypes, using the 2020 UKB imaging data release of approximately 40,000 subjects. The discovery sample has now almost tripled (22,138), the number of phenotypes increased to 3,935 and the number of genetic variants with MAF≥1% increased to 10 million. For the first time, we include associations on the X chromosome, and several new classes of image derived phenotypes (primarily, more fine-grained subcortical volumes, and cortical grey-white intensity contrast). Previously we had found 148 replicated clusters of associations between genetic variants and imaging phenotypes; here we find 692 replicating clusters of associations, including 12 on the X chromosome. We describe some of the newly found associations, focussing particularly on the X chromosome and autosomal associations involving the new classes of image derived phenotypes. Our novel associations implicate pathways involved in the rare X-linked syndrome STAR (syndactyly, telecanthus and anogenital and renal malformations), Alzheimer’s disease and mitochondrial disorders. All summary statistics are openly available for interactive viewing and download on the “BIG40” open web server.

---

UK Biobank (UKB) is now approximately halfway through imaging 100,000 volunteers; the early-2020 release of brain imaging data contained data from almost 40,000 participants. This spans 6 brain MRI (magnetic resonance imaging) modalities, allowing the study of many different aspects of brain structure, function and connectivity. In conjunction with other data being recorded by UKB, which includes health outcomes, lifestyle, biophysical measures and genetics, UKB is a major resource for understanding the brain in human health and disease.

In Elliott et al. [2018], we presented genome-wide association studies (GWAS) of 3,144 brain imaging phenotypes, with a discovery sample of 8,428 subjects. At that point we identified 148 replicated clusters of associations between genetic variants and the phenotypes. We found links between IDPs and genes involved in: iron transport and storage, extracellular matrix and epidermal growth factor, development, pathway signalling and plasticity.

We have now expanded and enhanced this work, with an almost threefold increase in sample size, an increase in the number of IDPs to almost 4,000, and with a focus on X chromosome associations [Wise et al., 2013] carried out for the first time. The new classes of IDPs, computed on behalf of UKB and released for general access, are: subnuclei volumes in amygdala, brainstem, hippocampus and thalamus, Brodmann area FreeSurfer metrics and FreeSurfer-derived white-grey intensity contrasts; Dale et al. 1999. We have also greatly expanded our set of imaging confound variables [Alfaro-Almagro et al., 2020], reducing the likelihood of finding artefactual associations. GWAS summary statistics and Manhattan plots for all 3,935 phenotypes are freely available for download from the Oxford Brain Imaging Genetics (*BIG40*) web server^1^, which also includes detailed tables of all IDPs, all SNPs (single nucleotide polymorphisms) tested, all association clusters, and an interactive viewer allowing for detailed interrogation of IDP associations with SNPs and nearby genes. We also provide a list of causal genetic variants for our top X chromosome clusters using a statistical fine-mapping approach [Hormozdiari et al., 2014].

We conducted sex-specific GWAS on autosomes (chromosomes 1-22) and the X chromosome, followed by a meta-analyses combining these, using Fisher’s method [Fisher, 1948]. The X chromosome accounts for about 5% of the human genome and incorporates over 1200 genes, including many which play a role in human cognition and development [Brenner, 2013]. However, testing for association with genetic variants on chromosome X requires special consideration [Clayton, 2008, Özbek et al., 2018, König et al., 2014]. While genetic females inherit two copies of the X chromosome, genetic males inherit only a single copy from the maternal line (here we refer to people with two X chromosomes as genetic females, and people with one X and one Y chromosome as genetic males). The short pseudoautosomal regions (PAR) on the ends of chromosome X are homologous with parts of chromosome Y and can be analysed in the same way as autosomal chromosomes. For the non-pseudoautosomal region, a mechanism has evolved to balance allele dosage differences between the genetic sexes (X chromosome inactivation, or XCI); during female development, one copy is randomly inactivated in each cell. This means that maternally and paternally inherited alleles would be expected to be expressed in different cell populations within the body approximately 50% of the time. However, this dosage compensation mechanism (DC) is imperfect; it is currently thought that only 60-75% of X-linked genes have one copy completely silenced in this way [Sidorenko et al., 2019].

To account for this in GWAS, it is common to follow Clayton [2008] and assume full dosage compensation: males are treated as homozygous females with genotypes coded (0,2) according to whether they have 0 or 1 copy of the alternative allele. In a joint analysis of females and males, simulation studies in Özbek et al. [2018] and König et al. [2014] suggest that type-I error control under this approach is reasonably robust to deviations from other assumptions (such as no sex-specific differences in allele frequencies), provided genetic sex is included as a covariate.

Recent studies have used the large sample sizes afforded by UKB to perform stratified analyses to estimate the degree of dosage compensation as a parameter across a broad variety of traits [Sidorenko et al., 2019, Lee et al., 2018]. These studies suggest that only a small proportion of genes escape XCI, although the appropriate amount of DC shows considerable variation amongst traits [Sidorenko et al., 2019]. For educational attainment, Lee et al. [2018] estimated a DC factor of 1.45, but concluded that little power was lost in their joint analysis irrespective of the assumed model.

While a joint analysis under a full DC model is a reasonable default, the power afforded by biobank-scale datasets also permits examination of possible sex-specific effects via stratified analyses. These stratified analyses can subsequently be meta-analysed [Sidorenko et al., 2019, Lee et al., 2018, Luciano et al., 2019]. If the meta-analysis is based on estimated effects (regression beta values), Lee et al. [2018] shows that appropriately chosen weights can give results almost the same as those from a joint genetic male/female analysis corresponding to any assumed DC model. Nevertheless, results will be biased if the assumed DC model differs from the truth. An unweighted meta-analysis based on p-values using Fisher’s method (explored in our research), though potentially less powerful, should avoid this possible bias (as it is not sensitive to any relative scaling in the regression model and hence the effect sizes in the two sex-separated GWAS), and still have value in confirming signals from a joint analysis.

The Oxford Brain Imaging Genetics (*BIG40*) web server includes a summary statistics resource including results for our discovery cohort, and a full GWAS on all samples passing QC (with discovery and replication cohorts combined). A browsable interface is also provided for both of these subsets. In addition, the results of the sex-separated GWAS on chromosome X and autosomes are provided. We have also provided the summary statistics to the European Bioinformatics Institute (EBI) GWAS repository. More details about our resource are provided in Appendix B of the Supplementary Material.

## Results

### Overview of GWAS Results

We conducted a genome-wide association study using the 39,691 brain imaged samples in UK Biobank. We divided these samples into a discovery (N=22,138) and a replication (N=11,086) cohort. The details for the imaging and genetics processing and the cohorts are given in Online Methods. We applied automated methods for identifying local peak associations for each phenotype, and also for aggregating peaks across phenotypes into clusters of association (as described in Appendix A of the Supplementary Material). A cluster is a set of phenotype/variant pairs such that all of the phenotype/variant pairs have a −Log10(*P*) value for association that exceeds a 7.5 genome-wide significance threshold, and such that all of the pairs have variants that are close with respect to genetic distance. We assigned each of the phenotype/variant pairs to one and only one cluster. We defined a cluster as replicating if at least one of the phenotype/variant pairs had nominal significance in our replication cohort (*p*-value < 0.05).

With these methods, we found 10,889 peak associations among all phenotypes and chromosomes (8,446 replicating at nominal significance), and found 1,282 clusters (692 replicating) after clustering the peak associations according to our automated methods (the number of replicating clusters reported in Elliott et al. 2018 was 148). The 692 replicating clusters are distributed across all chromosomes, with between 8 and 60 clusters per chromosome. We grouped the IDPs into 17 categories (Supplementary Table 1). Of the replicated associations among these 692 clusters, 16 out of 17 categories are represented (the *task fMRI activation* category is the only category without at least one association). The number of associations per category ranges between 12 for the category *volume of white matter hyperintensities (lesions)* which consists of just one IDP [Griffanti et al., 2016], and 1,954 for the *regional and tissue volume* category. All of these associations are listed in Supplementary Table S4, and Manhattan plots along with quantile plots are provided on the *BIG40* open web server.

Of all of our clusters of associations, 38 are on the X chromosome (12 replicating), and 4 of the X chromosome clusters have a phenotype/variant pair with association significance exceeding the more stringent Bonferroni^2^ corrected level of −Log10(*P*) ≥ 11.1. These four clusters are investigated below. We also investigate five novel clusters among the autosomal chromosomes.

We provide a fine-mapping of the four X chromosome clusters using the CAVIAR software [Hormozdiari et al., 2014] with results described in Supplementary Table S3, considering regions within 250kbp of the lead associations for the clusters. For cluster 1, this resulted in a region containing 1,211 genetic variants. We ran the CAVIAR software [Hormozdiari et al., 2014] with default settings and recorded the genetic variant found to be most causal for each phenotype association within the cluster. For cluster 1, CAVIAR reported rs2272737 (the same lead genetic variant found by our aggregation method) as the causal genetic variant for 81/97 of the phenotype associations. The situation was similar for the other three clusters; in each case, the lead genetic variant found by our aggregation method was among the genetic variants found to be causal by CAVIAR. (The number of genetic variants included in the 500kb regions examined by CAVIAR in the remaining three clusters included 2,017, 731 and 935 genetic variants, respectively. The proportion of phenotype associations for which the lead genetic variant found by our aggregation method was also estimated to be the most probable causal variant by CAVIAR for these remaining 3 clusters was 10/21, 5/35 and 5/17, respectively.) The CAVIAR results are detailed in Supplementary Table S3 and further details provided in the Supplementary Material.

BIG40 also provides summary statistics for a GWAS with the discovery and replication cohorts combined (a *full-scan*). We also examined the heritability of each phenotype using linkage score regression [Bulik-Sullivan et al., 2015]. The heritability for each of the phenotype categories is summarized in Figure 3. With phenotypes for which the estimated *h*^2^ value was more than one standard error greater than 0, the estimated value of *h*^2^ ranged from 0.01 to 0.41. The highest estimated heritability was found in phenotypes involved in regional and tissue volumes (as in Elliott et al. 2018), cortical grey-white contrast, and the intra-cellular volume fraction (ICVF) diffusion tensor imaging measure.

### X Chromosome Results - Overview and Sex-specific Tests

Full details for the lead associations for the X chromosome clusters (including clusters that do not replicate) are provided in Supplementary Table S2. A summary of all of the peak associations included in the X chromosome clusters is provided in Supplementary Table S3, and the full results for peak associations on all chromosomes are provided in Supplementary Table S4. In these tables, clusters numbers are given in the first column, and clusters are ordered based on the chromosome number (in ascending order with the X chromosome first) and then by the −Log10(*P*) value of the lead association (in descending order). A summary of all replicating X chromosome clusters is provided in Table 1, and further details including genes and expression quantitative trait loci (eQTL) for the four Bonferroni-significant clusters are provided in Table 2. Figure 1 shows Manhattan plots for the lead associations in these 4 top X chromosome clusters, and these clusters are explored further below.

**Figure 1:**
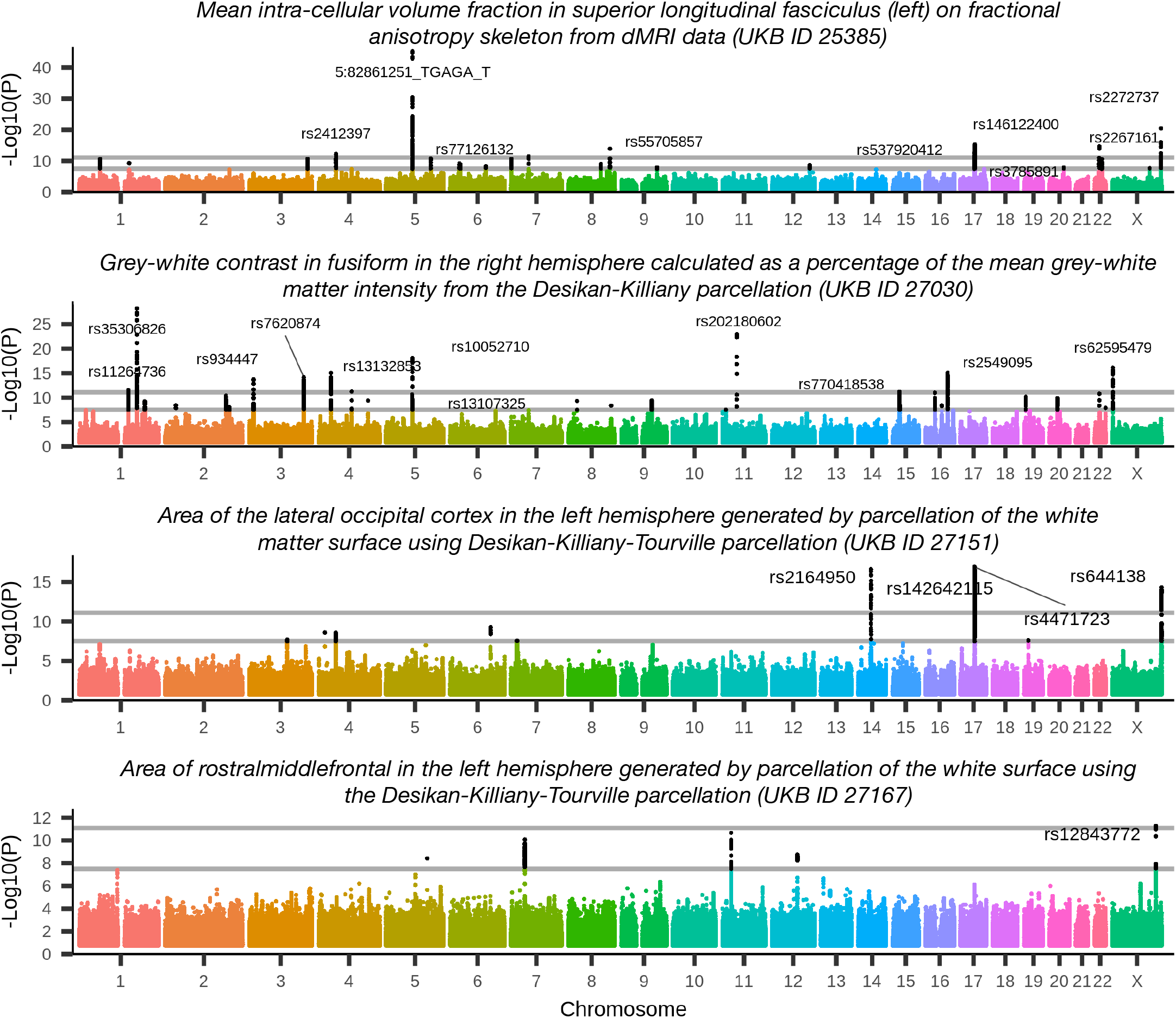
Manhattan plots for the four phenotypes achieving Bonferroni corrected significance on the X chromosome. Genetic variants are labelled for peak associations achieving the Bonferroni level. Plot titles indicate phenotype definition (including UKB ID field index from http://biobank.ctsu.ox.ac.uk/crystal/field.cgi?id=25385, 27030, 27151 or 27167). Black dots indicate associations that are significant associations at the genome-wide level, −Log10(*P*) ≥ 7.5. Grey lines show genome-wide+Bonferroni level (11.1) and genome-wide significance level (7.5). These associations involve diffusion MRI and the Desikan-Killiany and the Dessikan-Killiany-Tourville parcellations [Fischl et al., 2004] of white matter and grey matter.

**Figure 2:**
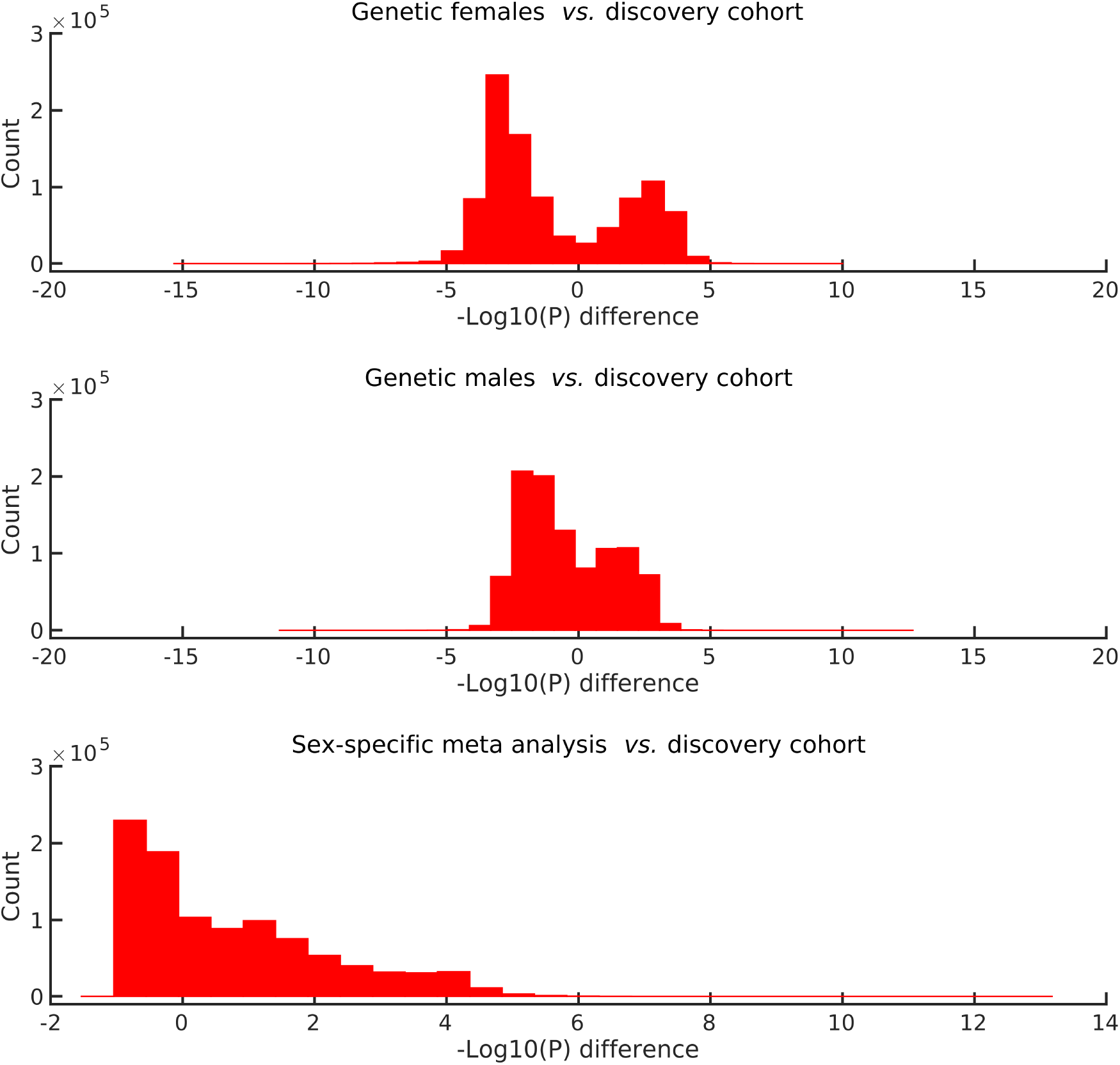
Paired difference histograms for the sex-specific scans on the X chromosome. We plot histograms for the differences between the −Log10(*P*) values for: genetic females (top), genetic males (middle), and the meta-analysis (bottom), vs. the discovery scan (which includes genetic male and female samples together, but did include a sex confound covariate). Differences are plotted for all associations for which the maximum −Log10(*P*) value over the four analyses is greater than 4.0, leading to the bimodal nature of the first two histograms. A total of 989,981 variants pass this maximum filter. The bottom plot shows that there is greater statistical sensitivity when carrying out sex-specific GWAS on the X chromosome, and then combining the results with a meta-analysis, than by combining all subjects together in a simple standard GWAS.

**Figure 3:**
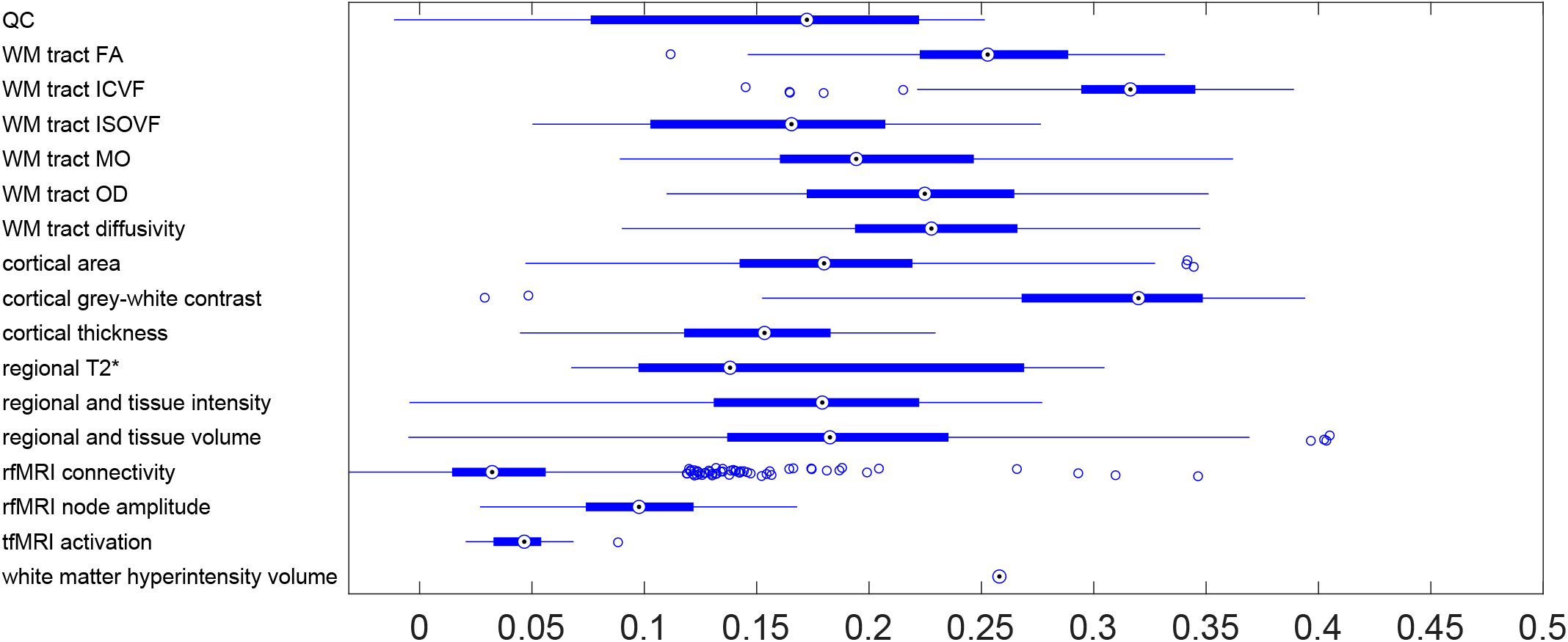
Heritability estimates (*h*^2^) for phenotypes grouped according to IDP categories. Acronymns in *y*-labels include: quality control (QC), and diffusion tensor imaging phenotypes: white matter (WM), fractional anisotropy (FA), intra-cellular volume fraction (ICVF), isotropic or free water volume fraction (ISOVF), diffusion tensor mode (MO) and orientation dispersion index (OD). More details for these 17 categories and heritability and standard errors for all phenotypes provided in Supplementary Table S1.

**Table 1:** Lead associations for the 12 replicating X chromosome clusters. Links to UKB phenotype definitions are provided in column 3. Summary statistics *beta* and *se* are provided in Supplementary Table S1. The −Log10(*P*) values provided are for the main discovery cohort. Column N_*SNPs*_ indicates the number of peak phenotype/variant pairs included in the cluster. Bolded rsids indicate significance at the Bonferroni threshold in the main discovery cohort.

| # | rsid | Phenotype ID and brief description | Position | a1 | a2 | −Log10(P) | N<sub>SNPs</sub> |
| --- | --- | --- | --- | --- | --- | --- | --- |
| 1 | **rs2272737** | 1943. White matter neurite density in left superior longitudinal fasciculus | 152876080 | C | T | 20.459 | 97 |
| 2 | **rs62595479** | 1408. White-grey T1 intensity contrast in right fusiform gyrus | 2163736 | T | C | 16.088 | 21 |
| 3 | **rs644138** | 0818. Area of left lateral-occipital cortex | 154927581 | G | A | 14.319 | 35 |
| 4 | **rs12843772** | 0834. Area of left rostral-middle-frontal cortex | 136648126 | C | T | 11.292 | 17 |
| 5 | rs188847578 | 0076. Volume of grey matter in left supplementary motor cortex | 127661277 | G | T | 10.247 | 3 |
| 10 | rs193203210 | 1437. Volume of white matter hyperintensities | 152634500 | C | G | 8.711 | 2 |
| 11 | rs11539157 | 0202. Volume of left ventral diencephalon | 68381264 | C | A | 8.659 | 1 |
| 15 | rs149776026 | 3512. Functional connectivity, connection 1084 dimensionality 100 | 124604066 | A | G | 8.193 | 1 |
| 16 | rs5916169 | 2510. Functional connectivity, connection 82 dimensionality 100 | 5507316 | C | T | 8.120 | 1 |
| 23 | rs6527976 | 1437. Volume of white matter hyperintensities | 13808841 | A | G | 7.981 | 1 |
| 25 | rs5930655 | 0136. Volume of grey matter in brain stem | 133781599 | A | C | 7.925 | 1 |
| 30 | rs6644158 | 0244. Volume of subiculum body in left hippocampus | 113638468 | G | A | 7.723 | 1 |

**Table 2:**
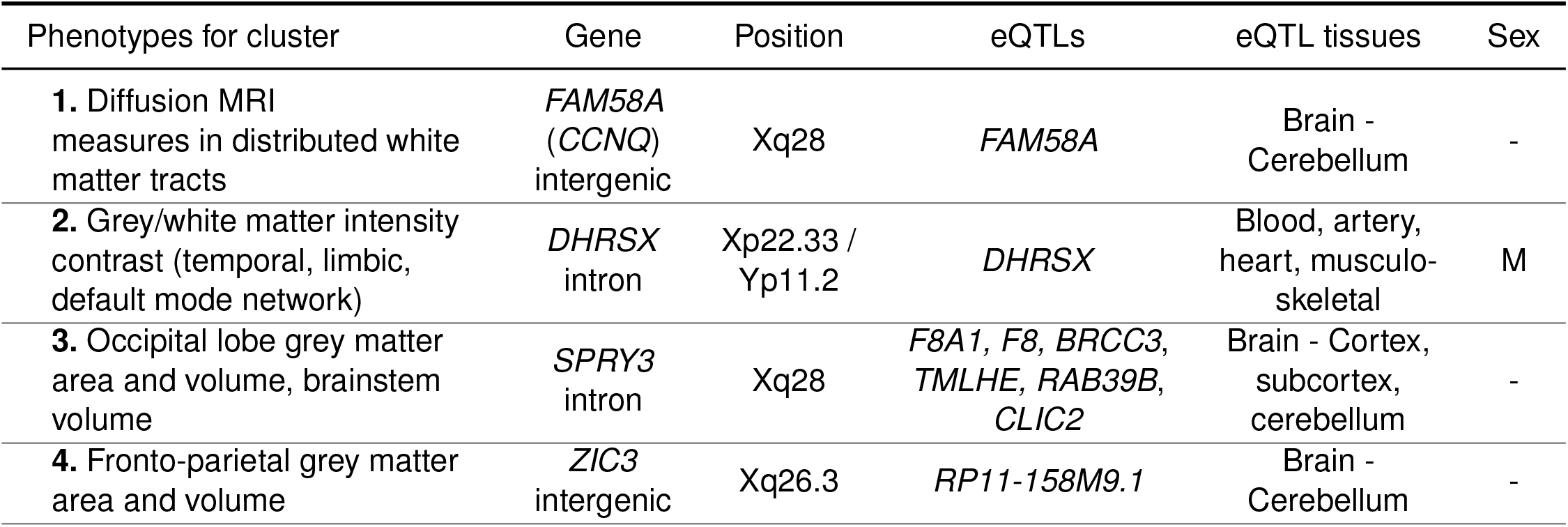
Context for the four X chromosome clusters with significance at the Bonferroni threshold. Cluster number (bolded and matching cluster numbering in the Supplementary Tables) and a general description of the phenotypes involved in the cluster are given in the first column. Cytogenetic positions are provided: clusters with both X and Y positions are in a pseudoautosomal region. Note that the cytogenic position Xq28 is on the edge of the non-PAR region and overlaps with the Y chromosome Yq12, although the lead association is in non-PAR. Information about expression quantitative trait loci (eQTLs) is provided. Column *sex* indicates if the sex-specific scan is significant at the Bonferroni threshold of −Log10(*P*) ≥ 11.1 for one genetic sex, but not significant for the other genetic sex (with *M* indicating the genetic sex for sex-driven effect in males, and dash indicating no such sex-driven effect).

Genetic sex affects the brain in fundamental ways [Nugent and McCarthy, 2011, Ruigrok et al., 2014, Saleem and Rizvi, 2017, Nguyen et al., 2019]. Our main GWAS analyses include sex as one of the confound variables (in part, as sex is a causal factor in some strong imaging confound effects such as interaction of head size with image intensity and head motion). To assess the quality of this deconfounding, and to explore associations on the X chromosome that are driven by genetic sex, we conducted two additional GWAS in which we restricted our discovery cohort to just genetic females and (separately) just genetic males. We then combined these two additional GWAS in a meta-analysis using Fisher’s method [Fisher, 1948]. Clusters in our main analysis that are significant in the meta-analysis but not significant in one of the sex-specific scans may indicate sex-driven associations. Of the 12 replicating X chromosome clusters in the main analysis (with genetic male and female sexes combined in the discovery cohort), one cluster (Cluster 2) is significant at the 11.1 level for one genetic sex, but not significant for the other genetic sex; Cluster 2 may therefore be driven by genetic males. To provide more direct evidence for this, we performed two-tailed z-tests to determine if the beta coefficients differ significantly between the genetic sexes [Clogg et al., 1995]. For Clusters 1 and 2, we found that the beta coefficients are nominally different (Cluster 1: P=1.2×10^−2^, beta coefficient for genetic females: −0.14, beta coefficient for genetic males: −0.08; Cluster 2: P=1.0× 10^−2^, beta coefficient for genetic females: 0.07, beta coefficient for genetic males: 0.13). The differences between the sex-specific beta coefficients for the lead associations of Clusters 3 and 4 were not significant. Among all of our associations (from all chromosomes and all IDPs) with at least one of the sex-specific scans significant at the −Log10(*P*) ≥ 11.1 level, the signs of the effect sizes for genetic females and genetic males always matched. For significance at −Log10(*P*) ≥ 7.5, the signs matched 99.42% of the time (Supplementary Figure S1).

Finally, we created an additional set of clusters (using the clustering method described in the Supplementary Material), based on the *p*-values of the meta-analysis of the X chromosome (thresholding at the genome-wide significance level). The clustering of the meta-analysis X chromosome scan produced 23 clusters. Twenty of these had lead associations within 0.25cM of one of the discovery cohort clusters derived from the original GWAS which pooled all subjects of both genetic sexes (indicating strong concordance between the meta-analysis and the discovery cohort). Each of the four discovery cohort X chromosome clusters with lead association at the Bonferroni level (the first four rows of Table 1) overlap with a meta-analysis cluster, suggesting that these main clusters are not confounded by genetic sex. Of the remaining meta-analysis clusters, three do not overlap with any of the discovery cohort clusters. The lead-rsid/lead-phenotype pairs of these three non-overlapping clusters are rs5990961/V3742, rs142994659/V1233 and rs764953454/V3919 (the mapping between the phenotype numbers and phenotype names is given in Supplementary Table S1). However, none of these three non-overlapping clusters achieved Bonferroni significance in the meta-analysis.

The differences in sensitivity between the main GWAS (pooling genetic male and female samples in the discovery cohort), the sex-specific GWAS and the Fisher meta-analysis are visualised concisely in Figure 2. The histograms show the distributions of paired-difference −Log10(*P*) values. For the sex-specific comparisons, there are SNP/phenotype pairs having reduced sensitivity compared with the original all-subjects GWAS (likely due to reduced statistical power because of reduced subject numbers), and other pairs with increased sensitivity (likely because a given association is stronger for the sex in question than for the other sex). The meta-analysis paired-difference distribution demonstrates that sex-separated GWAS followed by meta-analysis gives increased sensitivity to finding genetic associations in the X chromosome.

### Investigation of the Four Main X Chromosome Clusters

We now examine the four main X chromosome clusters in greater detail. The details for these additional investigations are summarized in Supplementary Table S5.

Cluster 1 comprises 5 SNPs, associated in total with 96 IDPs, all capturing differences in the properties of white matter tracts distributed throughout the cerebrum. This is described more in Table 2 and Supplementary Table S5. The top SNP (rs2272737, P=3.5 × 10^−21^) is located about 10 kb away from, and is an eQTL of *FAM58A* (or *CCNQ*). The eQTLs reported throughout this work were assessed as significant according to the Genotype-Tissue Expression (GTEx) project [GTEx Consortium, 2017]. Mutations in this gene lead to STAR (syndactyly, telecanthus and anogenital and renal malformations), a rare X-linked developmental disorder recently identified [Unger et al., 2008], for which notable brain variations have been observed such as incomplete hippocampal inversion, thin corpus callosum, ventriculomegaly and cerebellar hypoplasia [Bedeschi et al., 2017, Orge et al., 2016]. In addition, while *FAM58A* codes for an orphan cyclin with undescribed function, it has been shown recently to interact with *CDK10* [Guen et al., 2013]. Of particular relevance considering the many white matter IDPs associated with Cluster 1, mutation of the gene *CDK10* has been observed in a case study to lead to a rudimentary corpus callosum and paucity of white matter surrounding the lateral ventricles [Guen et al., 2018].

The SNPs of Cluster 1 are further associated with an array of non-imaging-derived phenotypes (nIDPs— phenotypes not derived from MRI) largely related to health (including diagnosed diseases and operative procedures), as well as some variables not necessarily health-related (Supplementary Table S5). Interestingly, one SNP in Cluster 1 (rs1894299) was seen previously in a GWAS of Type 2 diabetes [Suzuki et al., 2019]. This SNP is located in an intron of *DUSP9* (*MKP4*), a gene that codes for a phosphatase whose overexpression specifically protects against stress-induced insulin resistance [Emanuelli et al., 2008]. This may be related to another consistent aspect of these nIDP associations with Cluster 1: the diet of UK Biobank participants with intake of sweet food and drinks (including desserts, puddings, beer and cider).

Cluster 2 comprises 9 SNPs associated altogether with 17 IDPs, all of which are grey matter vs white matter intensity contrast, in limbic and temporal regions, and brain areas making up the default-mode network [Raichle et al., 2001]. The top SNP (rs62595479, P=8.2×10^−17^) is located in a pseudoautosomal region of chromosome X, i.e., a genetic region homologous between chromosomes X and Y — in an intron of *DHRSX*, and is an eQTL of the same gene. The genetic association with the grey-white intensity contrast IDP for this SNP was mainly driven by the male UK participants (as described above). The male-dominated aspect of the association between *DHRSX* and the brain has also been observed in a study showing that four PAR genes, including *DHRSX* (and *SPRY3*, see below), are up-regulated in the blood of genetic male patients with ischemic stroke [Tian et al., 2012].

While the majority of the nIDPs associations with the SNPs of Cluster 2 were related to diagnoses and operative procedures, half of these pointed at thyroid-related issues, in addition to the nIDP of workplace temperature, which may be related to thyroid function (Supplementary Table S5). Remarkably, the distribution of thyroid function modulation in the brain appears to consistently follow (in both positron emission tomography and fMRI studies), that of the 17 regions associated with Cluster 2: mainly limbic and temporal areas including the posterior cingulate cortex, orbitofrontal cortex, parahippocampal and fusiform gyrus [Miao et al., 2011, Schreckenberger et al., 2006, Zhang et al., 2014, Göttlich et al., 2015].

Cluster 3 includes 9 genetic variants associated with 28 IDPs of local brain volume. All 28 IDPs are located in the occipital lobe except for the volume of the brainstem and fourth ventricle. The top genetic variant (rs644138, P=4.8×10^−15^) is located in a PAR in an intron of *SPRY3*. This genetic variant is also an eQTL in many brain regions of a variety of genes whose mutations are involved in developmental and neurodevelopmental disorders: *RAB39B*, which plays a role in normal neuronal development and dendritic process, is associated with cognitive impairment [Vanmarsenille et al., 2014], X-linked intellectual disability [Giannandrea et al., 2010] and Waisman syndrome in particular, an X-linked neurologic disorder characterised by delayed psychomotor development, impaired intellectual development, and early-onset Parkinson’s disease [Wilson et al., 2014]; *TMLHE* is associated with X-linked autism [Celestino-Soper et al., 2011]; *CLIC2*, with X-linked intellectual disability [Takano et al., 2012]; and *BRCC3*, with an X-linked recessive syndromic form of moyamoya disease [Miskinyte et al., 2011]. It is also an eQTL of *F8*, and *F8A1* (*DXS522E/HAP40*), a likely candidate for the aberrant nuclear localisation of mutant huntingtin in Huntington’s disease. Considering the distribution of the brain IDPs in the occipital lobe, it perhaps lends additional credence to the consistent, but yet not understood, observation of volumetric and sulcal differences in these visual grey matter regions in Huntington’s disease gene carriers [Rosas et al., 2008, Mangin et al., 2020].

Another aspect of the genes for which the top genetic variant of Cluster 3 is an eQTL is cardiovascular issues: for instance, mutant *CLIC2* leads to atrial fibrillation, cardiomegaly and congestive heart failure [Takano et al., 2012], *F8* encodes a large plasma glycoprotein that functions as a blood coagulation factor, whereas mutations in *BRCC3* are linked to moyamoya syndrome, a rare blood vessel disorder in which certain arteries in the brain are blocked or constricted, and that is accompanied by other symptoms including hypertension, dilated cardiomyopathy and premature coronary heart disease [Miskinyte et al., 2011]. In line with this, we found that Cluster 3 was associated in UKB participants with diagnosis of atrioventricular block and ventricular premature depolarisation, as well as an operative procedure consisting of the replacement of two coronary arteries. In addition, Cluster 3 was consistently associated with many measures of physical growth, perhaps in line with the role of *SPRY3* as a fibroblast growth factor antagonist in vertebrate development, including height, lung function and capacity and body mass (Supplementary Table S5).

Finally, Cluster 4 comprised 8 genetic variants associated with volume of grey matter regions in the dorsolateral prefrontal cortex and the lateral parietal cortex (supramarginal gyrus and opercular cortex). The top genetic variant in this cluster (rs12843772, P=5.1 ×10^−12^) is located just <150 basepairs from *ZIC3*, which plays a key role in body pattern formation and left-right asymmetry. Mutations in this gene are thought to be involved in 1% of heterotaxy (situs ambiguous and inversus) in humans [Ware et al., 2004]. This may explain why the distribution of the higher-order grey matter regions associated with Cluster 4 follows the pattern of the fronto-parietal networks, which are notoriously left-vs-right segregated [Witt et al., 2020]. In particular, the supramarginal gyrus is known to show the strongest asymmetries from an early developmental stage [Dubois et al., 2010], and to be connected by white matter tracts that share a genetic influence with human handedness [Wiberg et al., 2019]. *ZIC3* is also involved in neural tube development and closure, and mutations in this gene cause, in addition to neural tube defects and cerebellum hypoplasia, consistent histological brain alterations with abnormal laterality and axial patterning, including a disorganised cerebral cortex [Purandare et al., 2002].

Cluster 4 is also in particular strongly associated with IGF-1 levels in the blood of UK Biobank participants (Supplementary Table S5). IGF-1 in particular controls brain development, plasticity and repair [Dyer et al., 2016]. More recently, it has emerged as a risk factor for dementia and particularly Alzheimer’s disease [Westwood et al., 2014], as it is a major regulator of *Aβ* physiology, and controls *Aβ* clearance from the brain [Bates et al., 2009]. Zoomed-in views of these associations are displayed in Supplementary Figure S2.

With respect to XCI status of the genes discussed in this section, the gene *DHSRX* is located in PAR1 and is fully expressed in females [Carrel and Willard, 2005]. Both *SPRY3* and *ZIC3* are subject to full inactivation [Balaton et al., 2015].

### Investigation of Five Autosomal Clusters

Among the 692 replicating clusters with GWAS significant p-values associations reported here, 48 involved a lead association among the autosomes with −Log10(*P*) ≥ 11.1, replication (at nominal level), and which were distinct from the loci reported in Elliott et al. 2018 and Zhao et al. 2019a, 2019b (*i.e*., more than 0.5cM from a genetic variant reported in Elliott et al. 2018 or Zhao et al. 2019a, 2019b, and more than 1Mbp from a gene reported as associated in Zhao et al. 2019a or 2019b). Of these 48 clusters, 7 involved associations with phenotypes from among the 3 new classes of UKB brain imaging phenotypes analyzed here. Of these 7 clusters, 2 (Cluster 271 with lead variant rs1368575 and Cluster 1070 with lead association rs3814883) were not related to previous brain imaging literature, and were not intergenic and were without eQTLs. The remaining 5 clusters (Clusters 163, 270, 841, 1029 and 1067 from the Supplementary Material Table S6) are investigated below. We provide a tabulation of all autosomal lead associations for our clusters with a cross-reference to Elliott et al. 2018 and Zhao et al. 2019a, 2019b in Supplementary Table S6.

The first of these most novel clusters related to previous brain imaging work (Cluster 163) included 12 distinct replicated variants associated with 31 IDPs, all but two related to the contrast between the intensity of the white matter and grey matter (tissue contrast: TC) generated by FreeSurfer [Dale et al., 1999], widespread across multiple cortical ROIs [Salat et al., 2009]. The top variant for this cluster (rs3832092, P=1.9 × 10^−13^) is in an exon and an eQTL of *MARS2*, a mitochondrial methionyl-tRNA synthetase, and is associated in particular with parietal TC (precuneus, inferior and superior parietal cortex). Mutations of *MARS2* have been linked to spastic ataxia, as well as neurodevelopmental delay, and white matter abnormalities (leukoencephalopathy), cortical and cerebellar atrophy, as well as corpus callosum thinning [Thiffault et al., 2006, Webb et al., 2015]. This variant is further associated primarily with many nIDPs related to mass and body fat, as well as nIDPs involving allergy, sleep and red blood cell count and shape (according to Open Targets Genetics; Carvalho-Silva et al. 2019).

The second cluster (Cluster 270) encompassed 13 replicated variants which were predominantly associated with the TC of 48 different cortical regions across the whole cerebrum. Its top variant (rs9290432, P=2.4 × 10^−17^), associated mainly with temporal, parietal and prefrontal TC, is in an intron and eQTL of the gene *PLD1*. This gene codes for a phospholipid enzyme that has been shown to regulate the trafficking of the protein *β*APP [Cai et al., 2006], the precursor of the amyloid beta whose plaques are a critical factor in the pathogenesis of Alzheimer’s disease. Increased expression of *PLD1* has been noted in the brains of Alzheimer’s disease patients both in the hippocampus and temporal lobe [Krishnan et al., 2018], and *PLD1* is upregulated in the mitochondrial membrane from brains of patients with Alzheimer’s [Jin et al., 2006]. The top genetic variant of this cluster is in addition related to nIDPs of red blood cell count and shape [Carvalho-Silva et al., 2019].

The third cluster (Cluster 841) consisted of 8 replicated loci all associated solely with measures of TC in 15 mainly higher-order cortical areas. We find that the lead variant in this cluster (rs549893, P=1.2 × 10^−12^), is associated with TC in the superior temporal cortex, and is located in an intron of *HMBS*, and is an eQTL of both *HMBS* and *VPS11* in many brain tissues (cortical, subcortical and cerebellar). The gene *HMBS* codes for an enzyme of the biosynthetic pathway of haem production. As such, mutations in *HMBS* are known to be related to acute intermittent porphyria, an autosomal dominant defect in the biosynthesis of haem. Acute intermittent porphyria manifests itself mainly as a neurological disorder, involving the autonomous, central and peripheral nervous systems, with acute life-threatening neurologic attacks. Deep cerebral white matter myelination abnormalities [Solis et al., 2004], and axonal neuropathy [Lindberg et al., 1996], have been observed in patients and animal models, as well as deficiencies in mitochondrial complexes in *Hmbs* mutant brain, suggesting that mitochondrial energetic failure also plays an important role in the expression of the disease [Homedan et al., 2015]. In addition, mutations in *VPS11* in 5 patients have been suggested as leading to infantile onset leukoencephalopathy with brain white matter abnormalities, severe motor impairment, cortical blindness, intellectual disability, and seizures, as well as in a significant reduction in myelination following extensive neuronal death in the hindbrain and midbrain in an animal model [Zhang et al., 2016]. The variant rs549893 is associated with nIDPs of body mass index, body fat mass and body fat percentage [Carvalho-Silva et al., 2019].

The fourth genetic cluster (Cluster 1029) comprised 17 replicated loci related again predominantly to TC in 45 widespread cortical areas. The lead variant for this cluster (rs140648465, P=4.1 × 10^−16^) is associated with TC measures in the supramarginal, inferior and superior parietal cortex, and it is intergenic, and is an eQTL in cortical and subcortical tissues of *RMDN3/FAM82A2*, coding for a regulator of microtubule dynamics. This tether protein is involved in facilitating the lipid transfer by increasing the contact sites between the endoplasmic reticulum and mitochondria, particularly in the brain [Fecher et al., 2019]. The protein FAM82A2 was found down-regulated in the frontal and parietal lobes in patients with Parkinson’s disease with dementia, with a dramatic reduction in the activity of the mitochondrial complexes [Garcia-Esparcia et al., 2018]. Recently, reduced levels of the RMDN3 protein (sometimes also named PTPIP51) have also been found in the temporal cortex of the brains of Alzheimer’s disease patients, suggesting that a disruption of endoplasmic reticulum-mitochondria interactions mediated by *RMDN3* might be part of the neuropathological process in Alzheimer’s [Lau et al., 2020]. A proxy genetic variant for the top variant of this fifth cluster (rs8042729, in LD of *R*^2^=1.00 with rs140648465) is further associated with nIDPs of blood count and hypertension [Carvalho-Silva et al., 2019].

The final interpretable cluster (Cluster 1067) comprised 31 replicated loci associated with a total of 51 IDPs of TC spread across higher-order cortical regions, as well as 19 other IDPs of cortical thickness (superior prefrontal and parietal), regional volumes (hippocampus, thalamus and choroid plexus) and white matter diffusion. This cluster’s most significant variant (rs2549095, P=5.3 × 10^−20^) is associated with TC in an array for temporal, parietal and prefrontal cortical areas. This locus is in an exon of the gene *CLEC18A*, which codes for a C-type lectin that has only been recently characterized, and shown to be expressed in the brain, in microglia [Huang et al., 2015]. This locus is also part of a 40-panel of genes that make it possible to distinguish cardioembolic from large-vessel ischemic stroke with high accuracy [Jickling and Sharp, 2015]. A proxy locus for the top genetic variant (rs72785089, LD *R*^2^=0.81) is further associated with measures of body mass and percentage, as well as blood count and alcohol intake frequency [Carvalho-Silva et al., 2019].

## Discussion

A major component in the expansion of the UK Biobank prospective epidemiological resource is the addition of tens of thousands of newly imaged participants, and the increase in the richness of phenotypes that can be derived from the imaging data. Since we published our first large-scale GWAS of UKB brain imaging in 2018, the brain imaging has almost reached its halfway point, having now scanned nearly 50,000 volunteers, of which 40,000 samples have already passed QC tests, and been processed and released for use in research. As a result, the size of the available discovery sample for GWAS has nearly tripled, and the number of genetic variants passing reliability thresholds (such as minor allele frequency MAF≥1%) has increased by 30%, now reaching 10 million. It is now therefore a good time to update our large-scale GWAS, now with almost 4,000 imaging-derived phenotypes - thousands of distinct measures of brain structure, function, connectivity and microstructure. The number of replicated clusters of imaging-genetic associations identified has more than quadupled since our previous work. We have made all of the GWAS summary statistics openly available via the new *BIG40* brain imaging genetics server.

We also studied brain imaging associations in the X chromosome, and autosomal associations with novel phenotypes. We identified 4 X chromosome clusters that replicate at the GWAS+Bonferroni level (increasing the standard 7.5 level according to Bonferroni correction for the number of IDPs tested). Among these four top X chromosome clusters, we find associations involving diffusion measures distributed in white matter tracts and grey/white matter intensity contrasts. We also find associations involving occipital lobe grey matter, and fronto-parietal grey matter. These associations are relevant to a diverse set of variations in brain development and pathologies including the recently identified STAR condition (syndactyly, telecanthus and anogenital and renal malformations), Waisman syndrome, early-onset Parkinson’s disease, X-linked autism, Alzheimer’s disease and Huntington’s disease. Cardiovascular conditions such as ischemic stroke, moyamoya syndrome and premature coronary heart disease are also implicated. The X chromosome genes *FAM58A (CCNQ), DHRSX, SPRY3, F8A1, F8, BRCC3, TMLHE, RAB39B, CLIC2* and *ZIC3* are involved in the associations we report.

The X chromosome is typically understudied in GWAS, and many of the X chromosome loci we identify are now implicated in genetic associations with brain imaging phenotypes for the first time. Ploidy, the Barr body and potential confounding with genetic sex complicates X chromosome analysis. To address this, we performed sex-specific GWAS, and a meta-analysis combining the sex-separated analyses. We showed significant overlap between the sex-separated meta-analysis and the main GWAS (the discovery cohort pooling genetic females and males), providing evidence against confounding (and we note the significant sex-specific effect signs never differed for genetic females and males for variants with MAF≥1%). This also allowed us to investigate if a given association was sex-driven (the association may be significant for only one genetic sex, or have significantly different effect sizes between the two sex-separated GWAS). For example, we found evidence that Cluster 2 for our brain-gene associations is driven by associations in genetic males.

Four of the autosomal clusters of association with grey-white tissue contrast were related to mitochondrial proteins and function. In the brain, mitochondrial disorders manifest themselves primarily as demyelinated lesions in the white matter, or as stroke-like lesions in both grey and white matter. More rarely, they present with cortical and subcortical necrosis [Finsterer, 2009]. While TC shows strong changes with ageing in prefrontal, lateral parietal, superior temporal and precuneus regions—those overall demonstrating the most significant associations with genetic variants—it is unclear what underlying tissue properties might be driving these effects in the MRI signal ratio [Salat et al., 2009]. Here TC, being almost entirely the only type of IDP showing significant genome-wide associations with our mitochondrial-related genetic clusters, appears to be specific and sensitive to mitochondrial function. These genetic-imaging findings are likely in line with the recent discovery that mitochondrial dysfunction, which emerges early during the ageing process, might prompt the catabolism of the myelin lipids of the white matter as an adaptive response to address brain fuel and energy demand [Klosinski et al., 2015]. This catabolic process perhaps explains the results previously observed with TC in ageing [Salat et al., 2009].

Enhancing our understanding of the mapping between genotype and phenotype may lead to advances in neuroscience and improvements in outcomes for brain pathologies. Deeply phenotyped resources such as UK Biobank provide an opportunity to update the known genotype/phenotype mappings for a wide spectrum of brain imaging phenotypes as more samples and more phenotypes are released. It is crucial that such updates are provided on open platforms, accelerating the potential impact of brain imaging genetics. We have done that here with the open *BIG40* resource (with a total of 1,282 clusters of associations), and with freely available summary statistics. We hope that this dissemination will be valuable for the next generation of neuroscience research.

## Acknowledgements

This research has been conducted in part using the UK Biobank Resource under Application Number 8107. We are grateful to UK Biobank for making the data available, and to all UK Biobank study participants, who generously donated their time to make this resource possible.

SS is supported by a Wellcome Trust Strategic Award 098369/Z/12/Z and a Wellcome Trust Collaborative Award 215573/Z/19/Z. LE and WC were funded by NSERC grants RGPIN/05484-2019, DGECR/00118-2019 and an NSERC Undergraduate Student Research Award. GD is supported by an MRC Career Development Fellowship (MR/K006673/1).

The *BIG40* Open Data Server is provided by The Wellcome Centre for Integrative Neuroimaging (WIN FMRIB), which is supported by centre funding from the Wellcome Trust (203139/Z/16/Z).

Compute resources were provided by the Oxford Biomedical Research Computing (BMRC) facility (a joint development between Oxford’s Wellcome Centre for Human Genetics and Big Data Institute, supported by Health Data Research UK and the NIHR Oxford Biomedical Research Centre). Some compute resources were also provided by ComputeCanada under the Resources for Research Groups program. We would like to thank the Resource Computing Managers at BMRC for their diligence in operation, with special thanks to Jon Diprose, Robert Esnouf and Adam Huffman. We would also like to thank Duncan Mortimer, the Senior Informatics Officer at BMRC, and Martin Siegert, the Research Computing Director and Site Lead at WestGrid/ComputeCanada. We are grateful to Sinan Shi, Anderson Winkler, Tom Nichols, Paul McCarthy, Doug Greve and Bruce Fischl for helpful input.

## Footnotes

1 Available at https://open.win.ox.ac.uk/ukbiobank/big40/

2 In this work we adjust for computing multiple GWAS (due to having multiple IDPs) by applying the (probably conservative) Bonferroni correction on top of the standard GWAS threshold of −Log10(*P*) ≥ 7.5, resulting in a ‘Bonferroni’ threshold of 11.1.

