## Supplementary Material for "Enhanced Brain Imaging Genetics in UK Biobank"

#### 1 Supplementary Methods

We describe the image processing and genetic preprocessing and the association studies, including the procedures that are X chromosome specific, and the details of our sex-specific analyses.

##### 1.1 Image Processing

We used brain IDPs from the "40k" (approximately 40,000 participants) UK Biobank data release in early 2020, as processed by WIN-FMRIB on behalf of UKB [Alfaro-Almagro et al., 2018]. After removal of subjects as part of the genetic processing (see below), we used data from 33,224 subjects. These were then randomly split into a discovery sample of 22,138 subjects (11,624 genetic females) and a replication sample of 11,086 subjects (5,787 genetic females). The ages in the discovery sample were: Females: mean age $= 63.6 \pm 7.3$ years, min $= 45.1$, max $= 81.8$. Males: mean $= 65.0 \pm 7.6$, min $= 46.1$, max $= 81.8$. In the replication sample: Females: mean $= 63.7 \pm 7.4$, min $= 46.3$, max $= 81.6$. Males mean $= 65.0 \pm 7.6$, min $= 46.1$, max $= 81.0$. The exact numbers of subjects vary across IDPs, according to patterns of missing data, with the maximum

numbers given above (for IDPs with no missing data), and the minimum numbers being just 16% lower. The BIG40 online table listing the IDPs includes the exact number of subjects (in discovery, replication samples and in the sex-specific GWAS) for each IDP. The details for these IDPs (including long descriptions, category names and units) are summarized in Supplementary Table S1.

As described in detail in Miller et al. [2016], the UK Biobank data includes 6 MRI modalities: T1-weighted and T2-weighted-FLAIR (Fluid-Attenuated Inversion Recovery) structural images, susceptibility-weighted MRI (swMRI), diffusion MRI (dMRI), task functional MRI (tfMRI) and resting-state functional MRI (rfMRI). We (and colleagues) have developed and applied an automated image processing pipeline on behalf of UK Biobank [Alfaro-Almagro et al., 2018], `https://www.fmrib.ox.ac.uk/ukbiobank/fbp`. This removes artefacts and renders images comparable across modalities and participants; it also generates thousands of image-derived phenotypes (IDPs): distinct measures of brain structure and function.

In this work we used the 3,913 IDPs available from UK Biobank, spanning a range of structural, diffusion and fMRI summary measures (described in the central UK Biobank brain imaging documentation `http://biobank.ctsu.ox.ac.uk/showcase/showcase/docs/brain_mri.pdf` and listed in full on the BIG40 server `https://open.win.ox.ac.uk/ukbiobank/big40/`).

We also used 16 QC measures available from UKB, as well as 6 compact summary functional connectivity features (derived from the hundreds of individual connectivity features; Elliott et al. 2018). In this paper we refer to all of the above 3,935 measures together as "the IDPs." Each IDP's $Nsubjects \times 1$ data vector had outliers removed (set to missing, with outliers determined by being greater than 6 times the median absolute deviation from the median). We discarded subjects where 50 or more IDPs were missing (for any reason, which could be due to: data acquisition incompleteness; data quality problems as described in Alfaro-Almagro et al. 2018, or the above-described outlier removal).

The data was then split into discovery and replication samples, and the remaining steps below applied to each sample separately. Each IDP's data vector was quantile normalised Miller et al. [2016], resulting in it being Gaussian-distributed, with mean zero, standard deviation one. Confounds were removed from the data, in a manner similar to that carried out in Elliott et al. [2018], including the 40 population genetic principal components supplied by UK Biobank and with a greatly expanded new set of confounds [Alfaro-Almagro et al., 2020]. This includes confounds for: age, head size, sex, head motion during functional MRI, scanner table position, imaging centre and scan-date-related slow drifts. In order to maximise GWAS interpretability, we regress out all confounds listed and recommended in Alfaro-Almagro et al. 2020. For higher-order (nonlinear and interaction) confounds, we used the same set of thresholds for automatic selection of these higher-order confounds. Given the slightly different set of subjects used here (for example, enforcing overlap with the genetics data) compared with those in Alfaro-Almagro et al. 2020, this resulted in the 602 'maximal' set of confounds (reported in Alfaro-Almagro et al. 2020) being reduced here to 597 confounds.

#### 1.2 Genetic Associations

We consider the 488,377 samples included in the Spring 2018 release of the UK Biobank, and proceed with a preprocessing and discovery/reproduction paradigm similar to that described in Elliott et al. [2018]. Of the samples, 39,944 are included among the 41,016 samples in the UK Biobank for which IDPs are available, after the genotyping quality control procedures for sample removal specified in Bycroft et al. [2018]. We removed samples without recent UK ancestry as determined by the *in.white.British.ancestry.subset* variable in the file *ukb_sqc_v2.txt* provided in the meta-data for UK Biobank. This variable selects samples based on self reported ancestry and genetic principal component thresholds. We also remove subjects based on relatedness, forming a maximal unrelated subset using the procedures recommended in Bycroft et al. [2018]. This results in a maximally unrelated subset of 34,298 samples with recent UK ancestry and accepted genotyping and imaging quality control. We divided this set into a discovery and reproduction cohort according to a random 2/3 and 1/3 (respective) proportion, resulting in a discovery cohort of 22,865 samples and a reproduction cohort of 11,433 samples.

For the X chromosome, fewer samples were available after exclusions due to variation in karyotype or additional aspects of genotype quality control described in Bycroft, Nature 2018. This resulted in discovery cohorts of 22,853 (instead of 22,865 as listed above) for the non-pseudoautosomal region and 22,844 for the pseudoautosomal regions, and replication cohorts of 11,430 for the non-pseudoautosomal region and 11,426 for the pseudoautosomal regions (X chromosome aneuploidy was assessed using the *putative.sex.chromosome.aneuploidy* column of the genetic quality control file released by UKB; Bycroft, Nature 2018).

Next, we consider quality control filters for the minor allele frequency (MAF), information score (INFO; Marchini and Howie 2010) and Hardy-Weinberg equilibrium (HWE). We apply filters for MAF $\geq$ 0.001 and INFO $\geq$ 0.3 and HWE $-\text{Log10}(P) \leq 7$. For the X chromosome, we apply these filters using the *qctool* software version 2.0.1 and the *-infer-ploidy-from sex* flag. This implies that genetic males contribute half as much as genetic females towards MAF and INFO for the non-pseudoautosomal region of the X chromosome. Further, as has been recommended [König et al., 2014], for the X chromosome the HWE filters are applied after computing for genetic females only. After these filters, of the 93,095,645 genetic variants included in UK Biobank in chromosomes 1:22, 16,445,196 remain. And of the 3,963,707 genetic variants in the X chromosome, 657,883 remain (639,835 in the non-pseudoautosomal region and 18,048 in the pseudoautosomal regions).

#### 1.3 Genome-wide Association Studies

We performed linear association tests on the samples in the discovery cohort. We performed these tests between each of the 17,103,079 genetic variants and each of the 3,935 IDPs described above (10,119,893 of which have MAF $\geq 0.01$). The genotypes were provided by UKB, and details for imputation (including X chromosome imputation) and genetic principal component construction are provided in Bycroft et al. 2018. We used the *bgenie* software [Bycroft et al., 2018] to conduct the GWAS and record the effect sizes (beta), standard errors and $-\text{Log10}(P)$ values for the associations. The effect sizes were recorded *in the direction of the alternate allele*. The phenotypes were scaled to have unit variance after deconfounding. The variants on chromosomes 1:22 were not scaled. Therefore, an effect size of 1.0 indicates that each copy of the alternate allele generally confers an increase in the phenotype by one standard deviation. For the non-pseudoautosomal region of X, the dosages for genetic males were scaled by a factor of 2.0 so that they lie in the range [0, 2], respecting the Barr body.

We produced Manhattan plots for each of the 3,935 IDPs, plotting the $-\text{Log10}(P)$ value for each variant (these Manhattan plots are provided on the BIG40 open web server, along with quantile plots). For the Manhattan plots, we applied an additional filter of MAF $\geq$ 0.01 to all variants. A method for extracting hits (peak associations) from a genome-wide association study was developed in Elliott et al. [2018]. We applied that method to extract hits from the 3,935 scans with $-\text{Log10}(P)$ values exceeding the GWAS threshold of 7.5 (again, with the MAF $\geq$ 0.01 filter), and annotated the Manhattan plots with these hits (note that in our tables we also signified results passing a Bonferonni corrected threshold for $p$-values below $1.0\times10^{-7.5}$ divided by the number of IDPs: $\sim$11.1). For each of the hits, we conducted a replication analysis by performing a linear association test on the samples in the replication cohort and record the effect sizes, standard errors and $-\text{Log10}(P)$ values for the replication. We extended the method from Elliott et al. [2018] in order to automatically generate clusters. A cluster is a set of phenotype/variant pairs such that each variant in the cluster is a peak association for its corresponding phenotype, and such that all variants are within a 0.25cM (centimorgans) distance of the phenotype/variant pair with highest $-\text{Log10}(P)$ value in the cluster. Each phenotype/variant pair identified as a hit appears in one and only one cluster. The details of this clustering method are provided in Appendix A of this Supplementary Material. We provide a software package called *Peaks* implementing these clustering methods, and have released it under the open source BSD 2-clause license.

Other methods for extracting lead associations include Bayesian methods such as CAVIAR [Hormozdiari et al., 2014] for determination of causal genetic variants. We examined causal genetic variants in the top four X chromosome clusters by computing the linkage disequilibrium matrix using *plink2* [Chang et al., 2015] for all variants within 250kbp from the lead associations (our CAVIAR results are provided in Table S3 of this Supplementary Material and the results are summarized in the main text). Summary statistics for the associations of all genetic variants (with MAF $\geq$ 0.001) for the discovery cohort (as well as the sex-separated and meta-analyses GWAS, and a pooled discovery+replication GWAS - see below), are available for download on the BIG40 open web server.

#### 1.4 Full-Scan

We also provide for download on the BIG40 open web server the summary statistics for a version of this genome-wide association study conducted on the union of the discovery and replication cohorts considered in this study (i.e., a maximal subset of unrelated samples with recent UK ancestry, among *all* samples in the UKB 2020 release of approximately 40,000 brain imaged samples). The genetic variants considered in this scan are the same genetic variants passing our filters for the discovery cohort reported on in this paper (with MAF $\geq$ 0.001). The sample size of each association test (after considering missing phenotypes and X chromosome exclusions due to aneuploidy) are also provided (the maximum number of included samples over all phenotypes in this full scan is 33,224).

Using the summary statistics from this full-scan, we estimate the heritability of each phenotype. We used linkage disequilibrium score regression [Bulik-Sullivan et al., 2015] to produce these estimates. Linkage disequilibrium scores were sourced from the European population of the 1000 Genomes project [The 1000 Genomes Project Consortium]. Results are listed in Supplementary Table S1.

#### 1.5 Sex-specific Scans

We also considered a scan for association on genetic male samples only, and also a scan for association on genetic females only (in both cases, with samples from the discovery cohort). For these scans, in the non-pseudoautosomal X chromosome region, 11,885 genetic female samples were used, and 10,968 genetic male samples were used. For the pseudoautosomal regions, 11,882 genetic female samples were used and 10,962 genetic male samples were used. As in the main analysis, the number of samples per phenotype varies due to missingness (the sample sizes for each phenotype for genetic females and males are provided in Supplementary Table S1). We conducted these two scans for association between each deconfounded phenotype and each variant on the X chromosome and the autosome, and recorded the effect sizes (beta), standard errors and $-\text{Log10}(P)$ values. We then combined these two scans in a meta-analysis using Fisher's method [Fisher, 1948], providing a $-\text{Log10}(P)$ value which is more strongly controlled for sex-specific effects. The equation for the combined $p$-value under this meta-analysis method is as follows:

$$1 - f_{\chi^2}(-2(\log p_m + \log p_f), 4) \tag{1}$$

Here $f_{\chi^2}(\,\cdot\,, \nu)$ is the cumulative distribution function of a chi-squared random variable with $\nu$ degrees of freedom, and $p_f$ and $p_m$ are the $p$-values of the genetic female and genetic male scans, respectively.

#### Software used

The following software packages and servers were used throughout this work:

- *bgenie* v1.3, software for efficient genome-wide association studies on high dimensional phenotype data. `https://jmarchini.org/bgenie/` [Bycroft et al., 2018]
- *qctool* v1.4 and v2.0.1, software for preprocessing genetic data. `https://www.well.ox.ac.uk/~gav/qctool_v1/` and `https://www.well.ox.ac.uk/~gav/qctool_v2/`
- *Peaks* v1.0, novel software for extracting clusters from multi-phenotype genome-wide association studies. `https://github.com/wnfldchen/peaks`
- *PheWeb* v1.1.19, a webserver for browsing phenome-wide associations. `https://github.com/statgen/pheweb` [Gagliano Taliun et al., 2020]
- *BIG40* open web server for Brain Imaging Genetics. `https://open.win.ox.ac.uk/ukbiobank/pheweb/`
- *plink2* v2.0, alpha software for conducting genome-wide association studies and preprocessing of genetic data `https://www.cog-genomics.org/plink/2.0/` [Chang et al., 2015]
- *CAVIAR* v2.0, fine-mapping software for extracting causal variants from summary statistics. `http://genetics.cs.ucla.edu/caviar/` [Hormozdiari et al., 2014]
- *Open Targets Platform* an online web server for genome-wide association study. `https://genetics-app.netlify.app/` [Carvalho-Silva et al., 2019]
- *LDSC* v1.0.1, software for heritability analysis from summary statistics (linkage score regression). `https://github.com/bulik/ldsc/` [Bulik-Sullivan et al., 2015]
- The GTEx online resource. `https://gtexportal.org/home/` [GTEx Consortium, 2017]

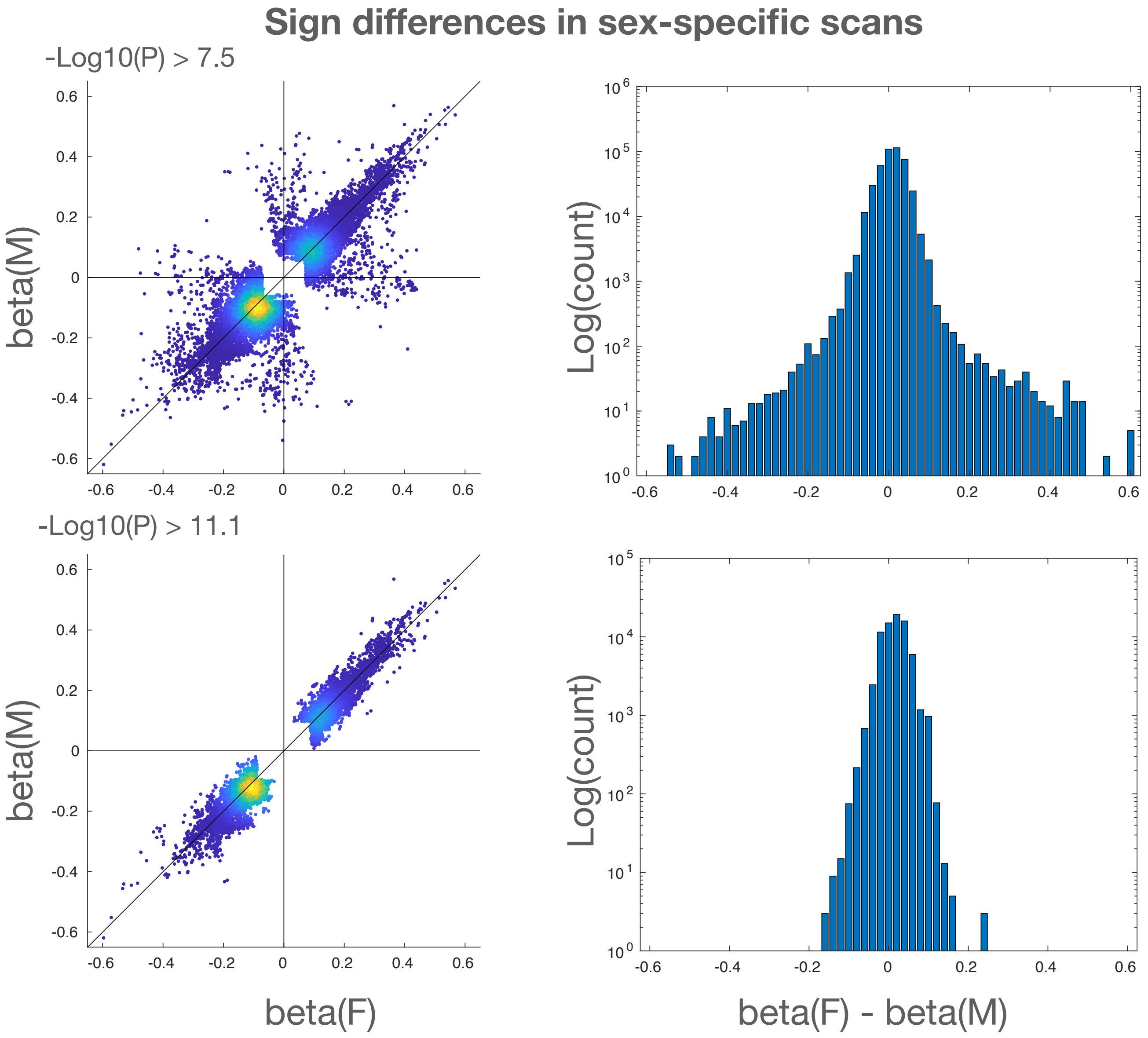


**Figure S1:** Comparison of signs for effect sizes for genetic females and males. *Top row*: Effect sizes for all associations with either genetic females or genetic males (or both) having $-\text{Log10}(P) \geq 7.5$. *Bottom row*: effect sizes for all associations with either genetic females or genetic males (or both) having $-\text{Log10}(P) \geq 11.1$. *Left column*: Scatter plots of effect sizes, indicating a small fraction (0.58%) of sign differences for $-\text{Log10}(P) \geq 7.5$ and no sign differences (quadrants II and IV empty) for $-\text{Log10}(P) \geq 11.1$ condition. *Right column*: Histograms of difference between effect sizes. Log $y$-scale indicates generally close matching of effect sizes.

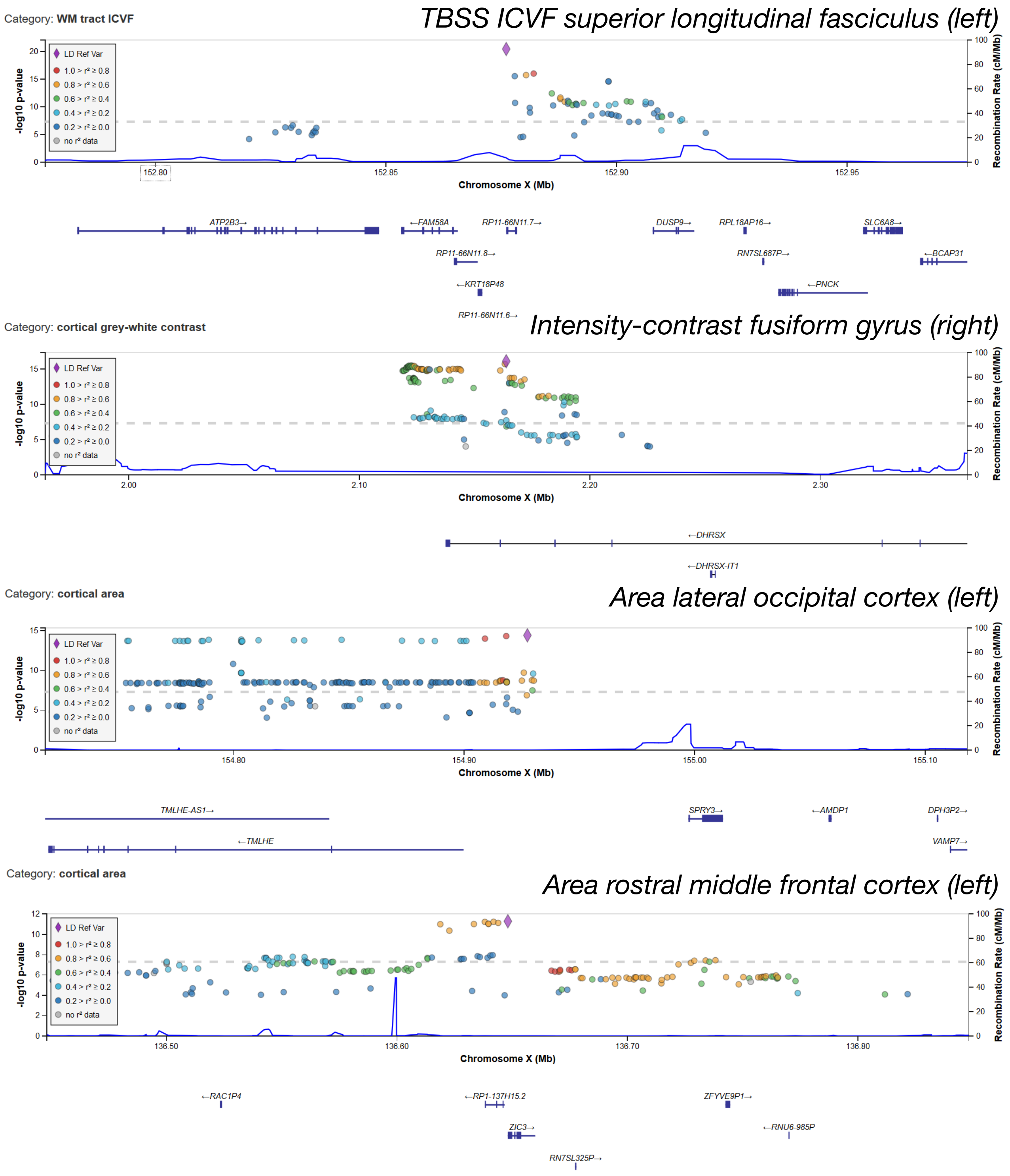


**Figure S2:** Regional association plots of the significant variants in X: (first row), loci within 10kbp of rs2272737 (P=$3.5 \times 10^{-21}$). And, an eQTL of *FAM58A*; (second row), rs62595479 (P=$8.2 \times 10^{-17}$) located in a pseudoautosomal region (PAR1) of X in an intron of *DHRSX*. And eQTL of the same gene; (third row), rs644138 (P=$4.8 \times 10^{-15}$) located in PAR2 in an intron of *SPRY3* (eQTL in brain tissue of various genes). Bottom row: rs12843772 (P=$5.1 \times 10^{-12}$) located $\leq$ 150bp from *ZIC3*. The genomic positions of the loci and genes are based on Human Genome build hg19.

### Appendix A: The *Peaks* algorithms

In genome-wide association studies with thousands of phenotypes, we must determine when two peak associations for two different phenotypes are related, in the sense that the peak variants are close together in genetic distance and/or linkage disequilibrium. For studies in which the number of phenotypes (or strength and complexity of genetic associations) amounts to only a few supra-threshold results, such matching can be done by hand by examining recombination maps and linkage-disequilibrium diagrams. Such by-hand work is not feasible for studies with thousands of phenotypes. In this work, we provide a new automated method to coregister peak associations across many phenotypes. We provide a software package *Peaks* which implements these methods.

#### Algorithm for Peak Associations

The *Peaks* software provide an implementation of the previously described algorithm in Elliott et al. [2018] (in the *Identifying associated genetic loci* subsection of the Methods section) for uncovering peak associations for each phenotype.

#### Algorithm for Cluster Identification

To combine peak associations into clusters that span phenotypes, we use a 'greedy' algorithm which delivers an optimally efficient clustering of genetic variant/phenotype pairs. The algorithm works by first identifying the peak associations for each phenotype (using the algorithm described in the previous subsection). Then, by iteratively extracting the genetic variant/phenotype pair with the top $-\text{Log}10(P)$, and then assigning all lead associations for all phenotypes within 0.25cM of that genetic variant to the same cluster. The details are as follows:

1. For each chromosome, convert the chromosome's peak associations into an array.
2. For each chromosome, convert the chromosome's array into a binary max-heap keyed on the $-\text{Log}10(P)$ of each genotype/phenotype pair, using the $\mathcal{O}(n)$ running time *heapify* algorithm described in Suchenek [2012].
3. For each chromosome, while the chromosome's heap is not empty, extract the maximum genetic variant/phenotype pair from the heap to create a new cluster. This can be done in $\mathcal{O}(\text{Log}(n))$ time. This extracted pair is the lead association of the cluster. Then, the 0.25 cM-cover is removed from the chromosome's heap, which is also an $\mathcal{O}(\text{Log}(n))$ operation. The cluster is outputted (including the removed aspects).
4. Since there are at most $n$ extractions and deletions the total running time is $\mathcal{O}(n\ \text{Log}(n))$, which is optimal as it is tight to the sorting lower bound of $\mathcal{O}(n\ \text{Log}(n))$, and since sorting is a relaxation of clustering and a lower bound of a relaxation has running time no worse than the original problem.

The open source *Peaks* software implementing these methods is available on *github* at `https://github.com/wnfldchen/peaks`.

### Appendix B: The BIG 40 online resource

Our resource includes openly released summary statistics and results for a variety of GWAS paradigms on the most recent release of 3,935 UKB brain imaging phenotypes. These results are released on BIG40 (`https://open.win.ox.ac.uk/ukbiobank/big40/`), the European Bioinformatics Institute (EBI) and the supplementary material of this paper. An enumeration of the aspects of our resource is as follows:

- Summary statistics for our disovery cohort ($N \leq$ ); available on BIG40 (Manhattan plots, full downloads, and a browsable interface), EBI under study accession numbers GCST90002426-6360 (`ftp://ftp.ebi.ac.uk/pub/databases/gwas/summary_statistics/GCST90002426` to `ftp://ftp.ebi.ac.uk/pub/databases/gwas/summary_statistics/GCST90006360`).
- Details of the clusters of associations identified by *Peaks*, including summary statistics for replication; available on BIG40 and in the Supplementary Material Tables S2 to S6.
- Causal variants identified by CAVIAR for the four X chromosome clusters significant at the $-$Log10($P$) $\geq 11.1$ level (Supplementary Table S5).
- A full GWAS on all phenotypes and chromosomes, with the discovery and replication cohorts combined (available on the BIG40 website as a download and as a browsable interface).
- Sex-specific GWAS on the discovery cohort with genetic females and genetic males considered separately, and combined through a Fisher meta-analysis (available as a download on BIG40).
- The heritability of each phenotype, assessed through LDSC on the full GWAS with discovery and replication cohorts combined (Supplementary Table S1).

The 1000 Genomes Project Consortium. A global reference for human genetic variation. *Nature*, 526(7571):68–74.
